# Bi-reporter vaccinia virus for tracking viral infections *in vitro* and *in vivo*

**DOI:** 10.1101/2021.08.24.457594

**Authors:** Kevin Chiem, Maria Lorenzo, Javier Rangel-Moreno, Maria De La Luz Garcia-Hernandez, Jun-Gyu Park, Aitor Nogales, Rafael Blasco, Luis Martínez-Sobrido

## Abstract

Recombinant viruses expressing reporter genes allow visualization and quantification of viral infections and can be used as valid surrogates to identify the presence of the virus in infected cells and animal models. However, one of the limitations of recombinant viruses expressing reporter genes is the use of either fluorescent or luciferase proteins that are used alternatively for different purposes. Vaccinia virus (VV) is widely used as a viral vector, including recombinant (r)VV singly expressing either fluorescent or luciferase reporter genes that are useful for specific purposes. In this report, we engineered two novel rVV stably expressing both fluorescent (Scarlet or GFP) and luciferase (Nluc) reporter genes from different loci in the viral genome. *In vitro*, these bi-reporter expressing rVV have similar growth kinetics and plaque phenotype than those of the parental WR VV isolate. *In vivo*, rVV Nluc/Scarlet and rVV Nluc/GFP effectively infected mice and were easily detected using *in vivo* imaging systems (IVIS) and *ex vivo* in the lungs from infected mice. We used these bi-reporter expressing rVV to assess viral pathogenesis, infiltration of immune cells in the lungs, and to directly identify the different subsets of cells infected by VV in the absence of antibody staining. Collectively, these rVV expressing two reporter genes open the feasibility to study the biology of viral infections *in vitro* and *in vivo*, including host-pathogen interactions and dynamics or tropism of viral infections. Moreover, they represent an excellent approach for the discovery of new prophylactics and/or therapeutics for the treatment of poxvirus infections.

**IMPORTANCE:** Despite the eradication of variola virus (VARV), the causative agent of smallpox, poxviruses still represent an important threat to human health due to their possible use as bioterrorism agents and the emergence of zoonotic poxvirus diseases. Recombinant vaccinia viruses (rVV) expressing easily traceable fluorescent or luciferase reporter genes have significantly contributed to the progress of poxvirus research. However, rVV expressing one marker gene have several constraints for *in vitro* and *in vivo* studies, since both fluorescent and luciferase proteins impose certain limitations for specific applications. To overcome these limitations, we generated optimized rVV stably expressing both fluorescent (Scarlet or GFP) and luciferase (Nluc) reporter genes to easily track viral infection *in vitro* and *in vivo*. This new generation of double reporter-expressing rVV represent an excellent option to study viral infection dynamics in cultured cells and validated animal models of infection, and for the discovery of new poxvirus antiviral treatments.

## INTRODUCTION

Poxviruses are large double stranded (ds)DNA viruses with a genome of ∼135-380 kb encoding up to 328 predicted open reading frames (ORFs) (1). The family *Poxviridae* includes several viruses of medical and veterinary importance. Some of the better-studied poxviruses belong to the Orthopoxvirus genus, which includes both vaccinia virus (VV), the prototypic member in the poxvirus family, and variola virus (VARV), the causative agent of smallpox. While smallpox has been eradicated, VARV still remains a pathogen of concern because of its potential use as a bioterrorism agent (2–4). Presently, much of the United States (US) population has not been vaccinated against smallpox (routine vaccination was discontinued in the 1970s) and, therefore, is susceptible to VARV infection (3, 5, 6). There is also a concern that monkeypox or additional Orthopoxviruses, can emerge and cause zoonotic disease due to a lack of pre-existing immunity within the human population (5).

VV constitutes a model virus for basic and biotechnological studies, and modification of its viral genome to express reporter genes has been vital in the study of viral gene expression, viral replication and pathogenesis, virus-host interactions, and cell entry mechanisms (7, 8). The development of reporter-expressing recombinant (r)VV has also been amendable to real-time and high-throughput screening (HTS) studies (9), and allow to easily assess viral infection without the use of laborious secondary methods to identify infected cells in culture or in live animals. However, to date, reporter-expressing rVV are limited to express a single reporter gene (e.g. fluorescence or luciferase proteins) or express distinctive fluorescent proteins using different promoters to monitor gene expression patterns (10, 11). This is typically because the development of reporter viruses has been largely motivated by a specific type of study, such as *in vitro* quantification of viral entry and replication, or by antiviral or neutralizing antibody (NAb) screening assays (9, 12, 13). Although these rVV expressing either fluorescent or luciferase proteins have proven extremely useful, they also have limitations based on the intrinsic properties of the reporter genes. Fluorescent proteins represent an excellent option for *in vitro* studies to track viral infection compared to luciferase proteins (14–17). However, luciferase proteins are more sensitive and convenient in quantitative analyses than fluorescent proteins (18–21). In live animals, luciferase activity can be easily tracked longitudinally using *in vivo* imaging systems (IVIS) and can be used as a valid surrogate of viral infection without the need of sacrificing animals (22). Contrary, fluorescent proteins represent a better option to identify the types of cells infected by the virus as they can be easily detected using fluorescent microscopy and flow cytometry assays (15, 23–25).

Here, we describe the generation of novel and stable rVV expressing both luciferase (Nluc) and fluorescence (Scarlet or GFP) reporter genes optimally designed for the easy detection of viral infection by both luminescence and fluorescence expression, respectively; and thereby, circumventing the limitations of using rVV expressing a single luminescence or fluorescent reporter gene. We selected Nluc based on its intense brightness, small size and ATP-independence (18, 23). We chose Scarlet or GFP, in combination with Nluc, to easily track viral infections based on the red or green fluorescent signal, respectively. The use of Scarlet (red) or GFP (green) fluorescent proteins would allow to track viral infections in genetically engineered cultured cells or animal models of infection. We selected GFP for its optimal brightness, and Scarlet for its excitation/emission at longer wavelengths, low toxicity and deeper light penetration in animal tissues (26). Notably, by introducing both fluorescent and bioluminescence reporter genes in the same virus, these rVV exploit the advantages of both fluorescent and bioluminescence proteins without the limitations of using one or the other.

*In vitro*, the plaque phenotype and growth kinetics of Nluc/Scarlet and Nluc/GFP bi-reporter rVV were comparable to the unmodified WR VV strain. Notably, reporter gene expression from the bi-reporter expressing rVV correlated with the levels of viral replication. *In vivo*, rVV Nluc/Scarlet and rVV Nluc/GFP were readily detected in real-time in infected animals by bioluminescence (Nluc), and *ex vivo* upon excision of lungs from infected animals (Scarlet or GFP) using an *in vivo* imaging system (IVIS). Furthermore, the spatial distribution of reporter viruses and their cellular targets in the lungs of infected mice were detectable in formalin-fixed, paraffin-embedded, tissue sections. Lastly, we determined the proportion of GFP-positive cells and visualized the spatial distribution of neutrophils in excised lungs from animals infected with rVV Nluc/GFP using flow cytometry and microscopy. Altogether, our studies indicate that these bi-reporter expressing rVV represent an excellent tool for studying the biology and immunology of VV *in vitro* and *in vivo*. The expression of two foreign genes inserted in different loci in the viral genome also opens the feasibility of generating new VV-based vaccine vectors expressing two foreign antigens for improving the treatment of other human pathogens. Notably, the generation of recombinant viruses expressing both fluorescent and bioluminescent proteins could be applied to other DNA viruses, allowing their study *in vitro* and *in vivo*, including the function of viral proteins, virus-host interactions, and the discovery of new antiviral therapeutic strategies.

## MATERIALS AND METHODS

### Cells

African green monkey kidney epithelial BSC-1 cells (ATCC CCL-26) were maintained in Eagle’s minimal essential medium (EMEM; Lonza, Inc.) containing 5% fetal bovine serum (FBS) and 1% PSG (100 U/ml penicillin, 100 µg/ml streptomycin and 2 mM L-glutamine) at 37°C with 5% CO_2_ supplementation.

### Viruses

Vaccinia virus (VV) Western Reserve (WR) strain was obtained from the American Type Culture Collection (ATCC VR-119) and routinely propagated in BSC-1 cells. Previously described vΔA27-ΔF13 is a VV mutant in which most of the coding sequence of genes A27L and F13L has been deleted (27).

### Plasmids

Plasmid pRB-NLuc, designed to express the NLuc gene downstream of the F13L gene has been previously described (27). Plasmid pA.S-TagGFP2 for insertion of GFP downstream of the A27L gene was constructed previously (27). A similar construct containing the gene coding for the red fluorescent Scarlet protein, derived from mCherry (28), was subcloned from plasmid pRB-Scarlet (27, 29) into the plasmid pA-LE-RGR (Sanchez-Puig and Blasco, unpublished), which contains A27L recombination sites flanking the A27L gene, to generate pA.S-Scarlet. For optimal expression, mScarlet was placed under a strong poxviral early-late synthetic promoter (30).

### Generation of recombinant vaccinia viruses (rVV)

rVV were generated as previously described (27). Briefly, BSC-1 cells (6-well plate format, 10^6^ cells/well), were infected at a multiplicity of infection (MOI) of 0.05 plaque forming units (PFU)/cell with vΔA27-ΔF13. After 1 h viral adsorption, virus inoculum was removed and cells were then transfected with 2 µg of plasmid DNA using FuGeneHD (Promega) according to the manufacturer’s recommendations. The rVV Nluc/GFP was generated by transfecting pRB-NLuc and pA.S-TagGFP2 plasmids. The rVV Nluc/Scarlet was obtained by transfecting pRB-NLuc and pA.S-Scarlet plasmids. After 2–3 days, cells were harvested, and rVV Nluc/GFP and rVV Nluc/Scarlet were isolated by three consecutive rounds of plaque purification. Viral stocks were generated in BSC-1 cells and viral titers were determined by standard plaque assay (PFU/ml).

### SDS-PAGE electrophoresis and Western blot analysis

Confluent monolayers of BSC-1 cells (6-well plate format, 10^6^ cells/well) were infected (MOI 0.05) with the indicated rVV. After 24 h infection, cell extracts were prepared in denaturant buffer (80 mM Tris-HCl, pH 6.8, 2% sodium dodecyl sulfate [SDS], 10% glycerol, 0.01% bromophenol blue solution and 0.71 M 2-mercaptoethanol). After SDS-polyacrylamide gel electrophoresis (PAGE), proteins were transferred to PVDF membranes and incubated 1 h at room temperature with primary antibodies diluted in PBS containing 0.05% Tween-20 and 1% nonfat dry milk. Primary antibodies include anti-NLuc rabbit polyclonal (1:6,000) (Promega), anti-mCherry rabbit polyclonal (1:5,000) (kindly provided by Dr. Antonio Alcamí), anti-GFP rabbit polyclonal (1:1,000) (Chemicon) rat monoclonal antibody 15B6, specific for F13 (1:50) (kindly provided by Dr. Gerhard Hiller), and anti-Actin monoclonal (1:500) (Sigma). After extensive washing with PBS containing 0.05% Tween-20, membranes were incubated with HRP-conjugated secondary antibodies diluted 1:3,000 in PBS containing 0.05% Tween-20 and 1% nonfat dry milk. Secondary antibodies were polyclonal goat anti-rabbit IgG (Dako), polyclonal goat anti-rat IgG (Dako) and polyclonal goat anti-mouse IgG (Sigma). After removal of unbound antibodies, membranes were incubated for 1 min with a 1:1 mix of solution A (2.5 mM luminol [Sigma], 0.4 mM ρ-coumaric acid [Sigma], 100 mM Tris-HCl, pH 8.5) and solution B (0.018% H2O2, 100 mM Tris-HCl, pH 8.5) to finally record the luminescence using a Molecular Imager Chemi Doc-XRS (Bio-Rad). The quantification of the bands was performed using the program Image Lab 3.0.1 (Bio-Rad).

### Plaque assay

To assess plaque phenotype of the rVV, BSC-1 cells (6-well plate format, 10^6^ cells/well) were infected with approximately 20 PFU per well. After 1 h infection, the medium was removed, and the infection was maintained for 48 h at 37°C under semisolid medium consisting of EMEM-2% FBS containing 1% low-melting point agarose. For fluorescence imaging, plates were photographed using the ChemiDoc™ XRS + (Bio-Rad). Green and red fluorescent signals from GFP and Scarlet, respectively, were photographed using the software settings indicated for SYBR Green using 1 s and 2 s exposure, respectively. For detection of Nluc, 1ml of PBS with 100 μl of substrate (Nano-Glo luciferase assay system, Promega) was added per well and immediately photographed with the software option for Chemiluminescence with 40 s of exposure. Cell monolayers were subsequently fixed with 1ml of 4% formaldehyde for 30 min at room temperature. After removing the semisolid medium, plaques were stained with a solution of 5% crystal violet in methanol (v/v) (31) and photographed using White Epi Illumination for 0.1 s.

### Viral growth kinetics

Viral growth kinetics were evaluated in BSC-1 cells (6-well plate format, 10^6^ cells/well, triplicates) infected at an MOI of 0.01. Viral replication at various times post-infection (24, 48, 72, and 96 h), was evaluated by phase contrast and fluorescence microscopy. In addition, at each time point, tissue culture supernatants were collected to determine viral titers and Nluc activity. Viral titers were determined by standard plaque assay (PFU/ml) followed by staining with crystal violet (31). Nluc activity was quantified using the Nano-Glo luciferase assay (Promega) following the manufacturer’s recommendations using an EnSight (Perkin Elmer) bioluminescence plate reader. Microsoft Excel software was used to determine the mean value and standard deviation (SD).

### Fluorescence microscopy

GFP and Scarlet fluorescence expression were detected using a Nikon Eclipse TE-2000-E inverted microscope. GFP expression was assessed using 465-495 nm (excitation) and 515-555 nm (emission) filters. Scarlet expression was detected using 515-560 nm (excitation) and 600-650 nm (emission) filters. Images were acquired with a Photometrics PRIME SCMOS monochrome camera.

### Mice experiment

All animal experiments were approved by the University of Rochester Institutional Biosafety (IBC) and Animal Care and Use (IACUC) Committees, which are in accordance with recommendations found in the Guide for the Care and Use of Laboratory Animals of the National Research Council (32). Six-to-eight-weeks-old female BALB/c mice were purchased from the National Cancer Institute (NCI) and maintained at the University of Rochester animal care facility under specific-pathogen-free conditions. Mice were anesthetized intraperitoneally (i.p.) with ketamine-xylazine (100 mg/kg ketamine and 10 mg/kg xylazine) and infected intranasally (i.n.) with the indicated viral PFU. Mice (n = 6/group) were daily monitored for changes in morbidity (body weight loss) and mortality (survival) for 14 days (33–35). Mice losing more than 25% of their initial body weight were considered to reach their end point and were humanely euthanized with carbon dioxide (CO_2_) and confirmed by cervical dislocation. The 50% mouse lethal dose (MLD_50_) was determined using the Reed and Muench method (36).

An IVIS Spectrum multispectral imaging instrument (Caliper Life Sciences, Inc.) was used for *in vivo* bioluminescence imaging of live mice and *ex vivo* imaging of lungs collected from mock-infected and VV infected animals. Six-to-eight-weeks-old female BALB/c mice (n = 4/group/day) were mock-infected (PBS) or infected i.n. with the indicated PFU. At the indicated days post-infection, mice were anesthetized with ketamine-xylazine (100 mg/kg ketamine and 10 mg/kg xylazine) and then retro-orbitally injected with 100 µl of Nano-Glo luciferase substrate (Promega) diluted 1:10 in PBS. Subsequently, mice were placed in an XIC-3 isolation chamber (Perkin Elmer) and imaged. Bioluminescence data collection and analysis were conducted using the Living Image software (v4.5; PerkinElmer). Flux measurements (Log_10_ photons per second (p/s) were obtained from the region of interest (ROI) around the whole body of each mouse. A bioluminescence intensity (radiance; number of photons per second per square centimeter per steradian; p/sec/cm^2^/sr) scale bar is displayed for each figure. Right after imaging, mice were removed from the XIC-3 isolation chamber and euthanized with a lethal dose of 2,2,2-tribromoethanol and exsanguination. Whole lungs were excised and washed with PBS. Scarlet or GFP expression were analyzed in the IVIS as previously described (15, 33, 37, 38). Living Image (v.4.5) software was used to acquire and analyze images to determine the radiant efficiency of the ROI. Fluorescence signals were normalized to those collected from mock-infected animals. Excised lungs from days 2, 4, and 6 post-infection were homogenized, and viral titers were determined by Median Tissue Culture Infectious Dose (TCID_50_).

### Histopathology and pathology scoring

Six-to-eight-weeks-old female BALB/c mice (n = 4) were infected (i.n.) with 10^7^ PFU of the indicated viruses. Mice were sacrificed on days 2 and 4 post-infection, and lungs were slowly perfused with 10% neutral formalin and fixed for 2 days. Next, the lungs were washed three times in PBS before being gradually dehydrated by sequential immersion in 70%, 95%, and absolute ethanol. After that, lungs were transferred to xylenes and embedded in paraffin. Lungs were sectioned (5 µm) and placed on a hot plate to dissolve the paraffin, followed by sequential rehydration with 95% ethanol, 75% ethanol, and ddH_2_O. Once the lungs were rehydrated, slides were mounted with Vectashield media containing DAPI (Vector Laboratories) to stain the nucleus. Lung sections were incubated overnight with Alexa Fluor 448 rabbit anti-GFP (Thermo Fisher Scientific), mouse anti-RFP (Thermo Fisher Scientific) or a rabbit polyclonal antibody against VV A33R protein (BEI Resources, NR-628) to demonstrate that fluorescent reporter proteins and viruses can be alternatively detected with antibodies. The following day, slides were incubated with rabbit anti-AF488 to amplify the GFP signal, Cy3 donkey anti-mouse IgG (Jackson ImmunoResearch Laboratories) to detect mouse monoclonal against Scarlet and with Alexa Fluor 647 donkey anti-rabbit IgG to detect the rVV. Slides were washed in PBS and mounted with Vectashield with DAPI (H-1200, Vector Laboratories). Representative pictures were taken with an Axioplan Zeiss Microscope and recorded with a Hamamatsu camera. Five randomly chosen independent anatomical locations from the lung sections were microscopically evaluated for histopathologic lesion scoring as previously described (39, 40). The histopathologic lesion scores included three different criteria (peri-bronchial & perivascular inflammation, intra-alveolar inflammation, and bronchial epithelial cell necrosis) and were graded on the basis of lesion severity as follows: grade 0 = no histopathological lesions; grade 1 = mild; grade 2 = moderate; grade 3 = marked; and, grade 4 = severe. A trained veterinary pathologist blindly performed all examinations at University of Rochester.

### Flow cytometry

Six-to-eight-weeks-old female BALB/c mice (n = 4) were infected (i.n.) with 10^7^ PFU of the indicated viruses and at days 2 and 4 post-infection, mice were sacrificed, and the lungs were surgically excised. Immediately afterwards, lungs were enzymatically digested with 0.625 mg/ml of collagenase (Sigma, C-7657) and 750 units of DNAse (Sigma) at 37°C for 30 minutes. Digested lungs were mechanically disrupted using metallic strainers. Cell suspensions were spun at 1,700 rpm for 4 min and red blood cells were lysed with ACK solution (0.15 M NH_4_Cl, 1 mM NaHCO3, 0.1 mM EDTA in PBS). Lived cells that excluded trypan blue were counted in a Neubauer chamber. Then, 50 μl of rat anti-CD16/CD32 (Fc block, clone 2.4 G2, Bioxcell, Lebanon NH), at 1 μg/ml, was added to the cell suspensions to prevent non-specific binding of fluorescently labeled antibodies. Cells were incubated for 15 min on ice with the following antibodies; Pe-Cy7 rat-anti-mouse CD45 (Clone 30-F11, Biolegend), PerCP-Cy5.5-rat anti-mouse IA-IE (Clone M5/114.15.2, Biolegend), APC mouse anti-mouse CD64 (Clone X54-5/7.1, Biolegend, San Diego, CA), APC-Cy7 rat anti-mouse CD11b (Clone M1/70, Biolegend, San Diego, CA), BV421 Armenian hamster anti-mouse CD11c (Clone N418, Biolegend, San Diego, CA), PE rat anti-mouse Ly6C (Clone HK1.4, Biolegend, San Diego, CA) AF700 rat anti-mouse Ly6G (Clone 1A8, Biolegend, San Diego, CA), biotin rat anti-mouse Siglec F (Clone S17007L, Biolegend, San Diego, CA). After incubation, cells were washed with FACS buffer (PBS containing 3% FBS and 2.5 mM of EDTA pH 7.4) and spun at 1,700 rpm for 4 min. Cells were incubated with Streptavidin PE-Cy5 (405205, Biolegend) for 10 min on ice, washed with FACS buffer, and then spun at 1,700 rpm for an additional 4 min. Lastly, cells were reconstituted with FACS buffer containing 1 μg/ml of propidium iodide and collected in an LSRII flow cytometer. Flow cytometric analysis was performed in the FlowJo software.

### Genetic stability of rVV

To test the genetic stability of rVV, viruses were serially passaged 5 times in BSC-1 cells (6-well plate format, 10^6^ cells/well) and incubated for 48 h until cytopathic effect (CPE) was observed. First passage was initiated using an MOI of 0.1 and the inoculum of the successive passages were 1:20 dilutions of the crude virus preparation from the previous passage. After 5 consecutive passages, viruses were monitored for the presence of fluorescence and luminescence using standard plaque assay. At least 50 plaques were counted for each virus.

### Statistical analysis

The One-way ANOVA or two-tailed paired/unpaired student’s T-test were used for statistical analysis on Graphpad Prism or Microsoft Word software, respectively; * p < 0.05, ** p < 0.01, *** p < 0.001, **** p < 0.0001 or no significance (n.s.)

## RESULTS

### Generation of rVV Nluc/Scarlet and rVV Nluc/GFP

The rVV Nluc/Scarlet and rVV Nluc/GFP were generated by insertion of Nluc downstream the F13L locus and either Scarlet (rVV Nluc/Scarlet) or GFP (rVV Nluc/GFP) downstream of the A27L locus (**Figure 1**). Insertion of foreign reporter genes was accomplished by a selection method based on plaque formation that allows simultaneous introduction of two genes into the VV genome (27). The corresponding genomic regions around those genes are depicted for the reference VV WR strain and for the recombinant viruses (**Figure 1A**). Notably, the insertion is directed into intergenic regions (F13L/F12L and A27L/A26L) and did not cause any additional modifications of the viral genome (**Figure 1A**).

**Figure 1.**
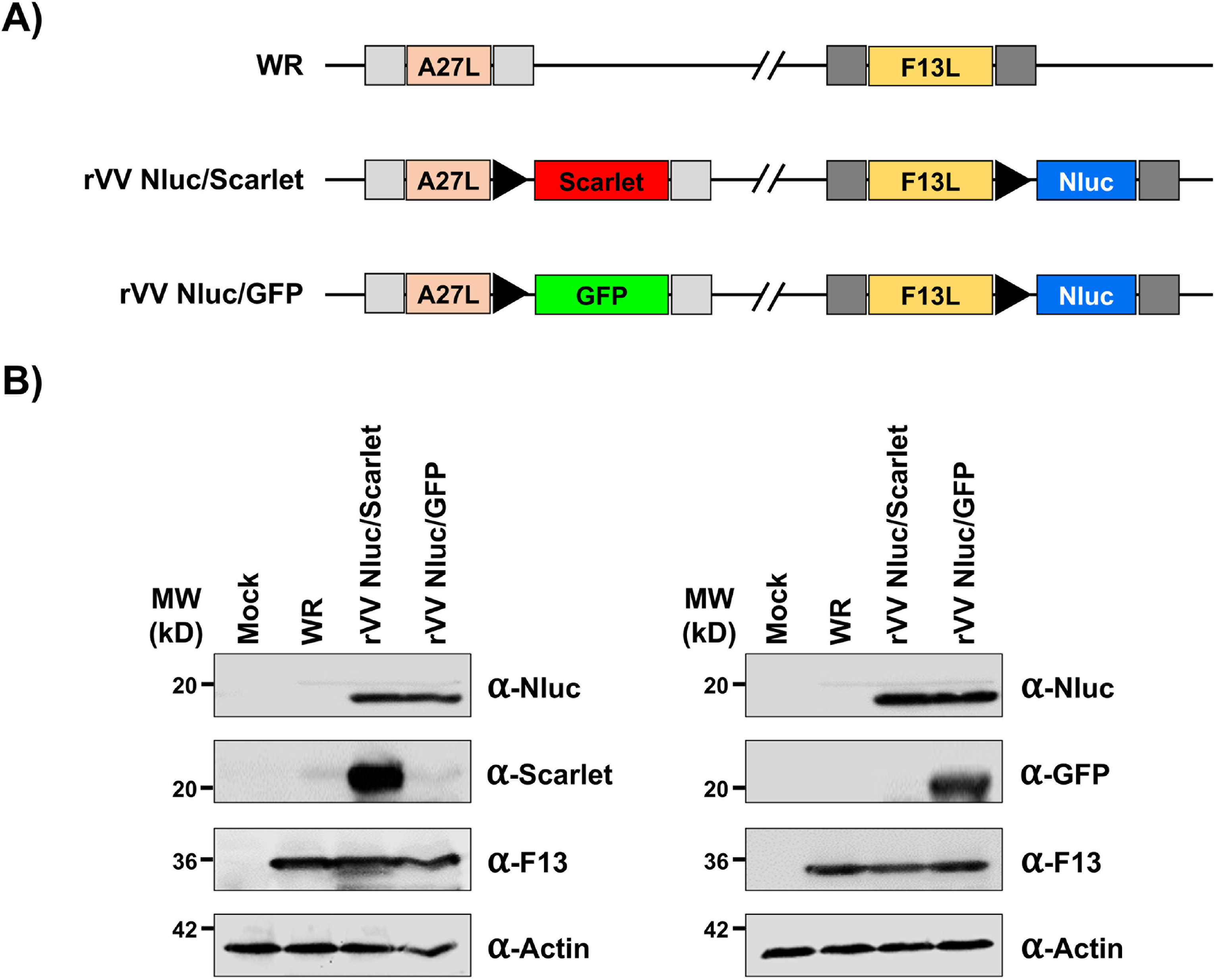
Generation of bi-reporter expressing rVV. **A) Schematic representation of WR (top), rVV Nluc/Scarlet (middle), and rVV Nluc/GFP (bottom) viral genomes:** Insertion sites downstream of the A27L (left) and F13L (right) genes in the viral genome are shown. Scarlet fluorescent protein (Scarlet; red), green fluorescent protein (GFP; green), and Nano luciferase (Nluc; blue) are shown. Fluorescent Scarlet and GFP were cloned downstream A27L (left). Nluc was cloned downstream F13L (right). Fluorescent Scarlet and GFP, and Nluc are expressed from an early/late VV synthetic promoter (black triangles). Gray boxes indicate the position of recombination flanks included in the plasmid to direct insertion in the VV genome. **B) Reporter gene expression:** Cell extracts from mock, WR, rVV Nluc/Scarlet and rVV Nluc/GFP infected (MOI 0.05) BSC-1 cells were collected at 24 h post-infection and analyzed for Nluc, Scarlet, and GFP expression by Western blot. An antibody against VV F13L and an antibody against actin were used as viral and cellular protein loading controls, respectively. The molecular size (kD) of the cellular/viral proteins are indicated on the left

We first characterized the rVV by Western blot (**Figure 1B**). Total cell lysates from mock-, WR-, rVV Nluc/Scarlet-, or rVV Nluc/GFP-infected (MOI 0.05) BSC-1 cells were collected at 24 h post-infection and they were analyzed using antibodies specific for Nluc, Scarlet (anti-mCherry), and GFP. As expected, Western blot analysis revealed specific bands with the predicted molecular size for Nluc only in cell extracts from BSC-1 cells infected with rVV Nluc/Scarlet or Nluc/GFP (**Figure 1B**). Furthermore, specific bands for Scarlet or GFP were only detected in cell extracts from rVV Nluc/Scarlet- or rVV Nluc/GFP-infected cells, respectively (**Figure 1B**). Antibodies against VV F13L and cellular actin were included as loading controls for viral infection and cellular housekeeping proteins, respectively (**Figure 1B**). As expected, we detected F13L only in VV-infected cell lysates, and the levels of F13L expression were similar between WR-, rVV Nluc/Scarlet-, and rVV Nluc/GFP-infected cells (**Figure 1B**). These results confirmed that each rVV expressed two reporter genes.

### *In vitro* characterization of reporter gene expression and viral replication of rVV Nluc/Scarlet and rVV Nluc/GFP

Next, we sought to assess whether the expression of reporter genes could be directly visualized by plaque assay and fluorescence microscopy (**Figure 2**). We conducted plaque assays on BSC-1 cell monolayers infected (∼20 PFU/well) with WR, rVV Nluc/Scarlet, or rVV Nluc/GFP. After 2 days post-infection, plaques were detected by either crystal violet staining, fluorescence (GFP and Scarlet), or bioluminescence (Nluc) (**Figure 2A**). Crystal violet staining revealed plaques with similar sizes for WR, rVV Nluc/Scarlet, and rVV Nluc/GFP (**Figure 2A**). As expected, only rVV Nluc/Scarlet and rVV Nluc/GFP, but not WR, displayed fluorescent positive plaques, respectively, under direct fluorescent imaging (**Figure 2A**). Moreover, in the presence of Nluc substrate, both rVV Nluc/Scarlet and rVV Nluc/GFP, but not WR plaques, were detectable using bioluminescence (**Figure 2A**). When representative images were merged, fluorescent plaques colocalized with Nluc-positive plaques, indicative of rVV Nluc/Scarlet and rVV Nluc/GFP expressing both reporter genes (**Figure 2A**). Moreover, Scarlet-, or GFP-, and Nluc-positive plaques colocalized with plaques detected by crystal violet, as indicated by red and green arrows, respectively (**Figure 2A**). Fluorescence expression of Scarlet and GFP in viral plaques from cells infected with rVV Nluc/Scarlet and rVV Nluc/GFP but not with WR, was further confirmed by imaging individual plaques under a fluorescence microscope (**Figure 2B**).

**Figure 2.**
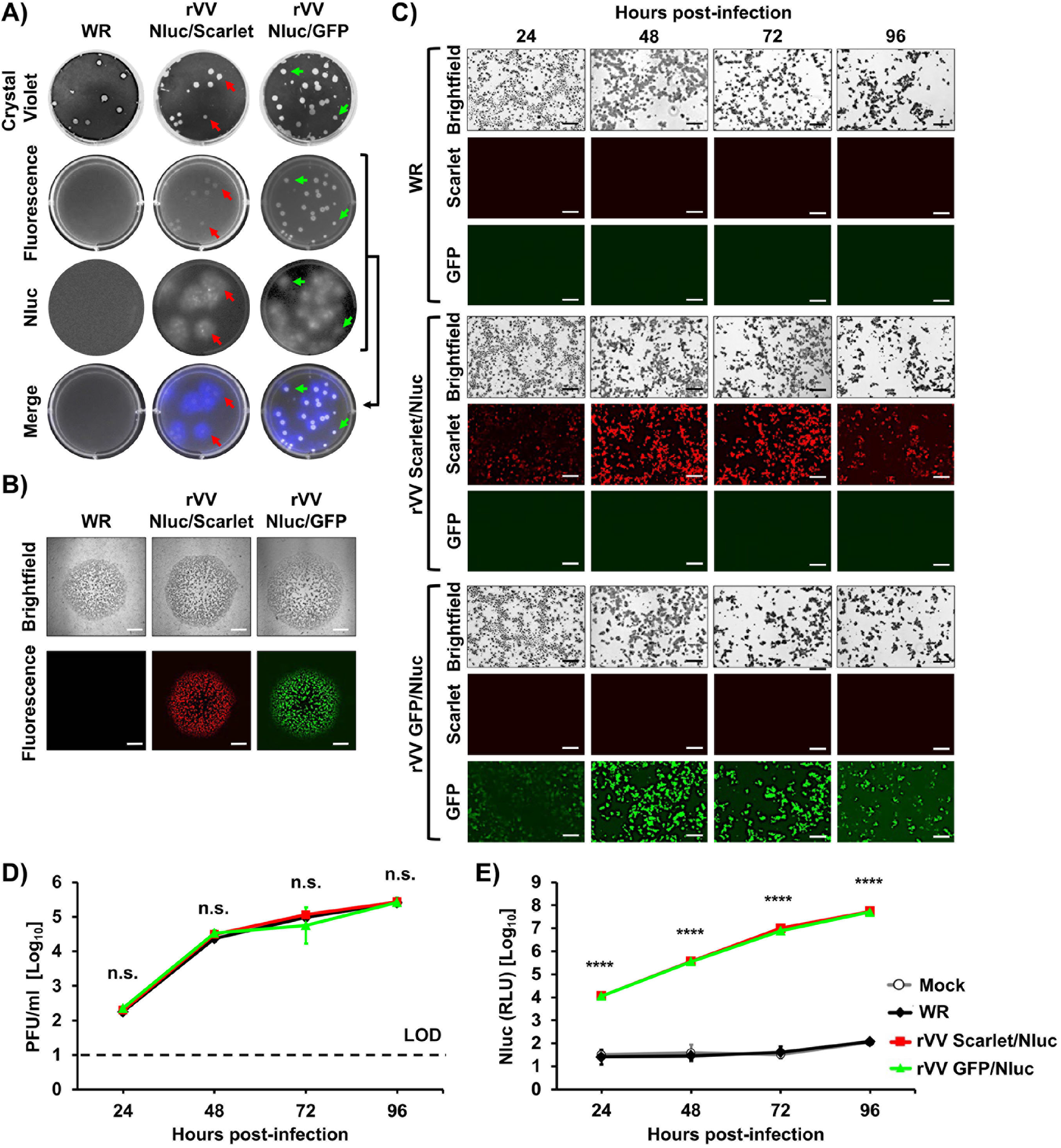
*In vitro* characterization of bi-reporter expressing rVV. **A) Plaque assay:** Representative images of WR (left), rVV Nluc/Scarlet (middle) and rVV Nluc/GFP (right) plaques in BSC-1 cells at 2 days post-infection are shown. Plaques were detected by crystal violet staining, Scarlet or GFP (fluorescence), and Nluc expression. Merged images are shown at the bottom. Red (rVV Nluc/Scarlet) and green (rVV Nluc/GFP) arrows show the superimposed images of fluorescence, respectively, and the crystal violet and Nluc-expressing plaques. **B) Fluorescence of individual plaques:** Plaques from BSC-1 cells infected with WR (left), rVV Nluc/Scarlet (middle) or rVV Nluc/GFP (right) were imaged using brightfield (top) or fluorescent (bottom) microscopy. **C) Fluorescent expression kinetics:** BSC-1 cells infected (MOI, 0.01) with WR (top), rVV Nluc/Scarlet (middle), or rVV Nluc/GFP (bottom) were observed using brightfield (top) and red (Scarlet, middle) and green (GFP, bottom) fluorescence microscopy at the indicated times post-infection. Representative images were taken at 200x magnification. Scale bars 0.5 mm. **D) Multicycle viral growth kinetics:** Tissue culture supernatants from BSC-1 cells infected (MOI, 0.01) with WR, rVV Nluc/Scarlet, or rVV Nluc/GFP were collected at the indicated times post-infection and assessed for the presence of virus by plaque assay. Data represent the mean and standard deviation of triplicates. The dotted line indicates the limit of detection (LOD, 10 PFU/ml). n.s: not significant differences. **E) Nluc expression:** Tissue culture supernatants from cells infected in panel D were used to determine levels of Nluc expression. RLU: relative light units. One-way ANOVA was used for statistical analysis; **** p < 0.0001.

To evaluate the kinetics of reporter gene expression, BSC-1 cells were infected (MOI, 0.1) with WR, rVV Nluc/Scarlet, or rVV Nluc/GFP and red or green fluorescence expression in infected cells was directly observed under a fluorescence microscope at 24, 48, 72, and 96 h post-infection (**Figure 2C**). Irrespective of the fluorescent protein used, signal was first detected after 24 h, progressively increased, and peaked at 72 h post-infection, with lower expression at 96 h post-infection, probably due to leakage from cells coincident with cytopathic effect (CPE) caused by viral infection (**Figure 2C**). By brightfield microscopy, WR-infected cells displayed viral-induced CPE similar to that of the rVV Nluc/Scarlet, or rVV Nluc/GFP but, as expected, fluorescence was only detected at background levels in WR-infected cells (**Figure 2C**).

Cell culture supernatants were also collected at the indicated time points to quantify viral titers (**Figure 2D**) and Nluc activity (**Figure 2E**). Viral titers in the tissue culture supernatants of BSC-1 cells infected with rVV Nluc/Scarlet or rVV Nluc/GFP were identical and comparable to those of WR-infected BSC-1 cells, with all viruses achieving similar viral titers at all times post-infection (**Figure 2D**). When we analyzed cell culture supernatants for Nluc activity, Nluc was detected as early as 24 h post-infection in BSC-1 cells infected with rVV Nluc/Scarlet or rVV Nluc/GFP (**Figure 2E**). Notably, Nluc expression levels increased in a time-dependent matter, peaking at 96 h post-infection, most likely because the CPE caused by viral infection resulted in the release and accumulation of NLuc in the tissue culture supernatants from infected cells (**Figure 2E**). As expected, BSC-1 cells infected with WR did not show detectable levels of Nluc in the tissue culture supernatant, like those of mock-infected cells (**Figure 2E**).

### Pathogenicity of bi-reporter expressing rVV in BALB/c mice

To be useful as a model for animal infection studies, it is critical that rVV retain the pathogenic potential of the unmodified virus. To assess whether rVV Nluc/Scarlet or rVV Nluc/GFP were pathogenic in mice, and to the level of WR, six-to-eight-week-old female BALB/c mice (n = 6/group) were inoculated i.n. with 10^4^, 10^5^, 10^6^, and 10^7^ PFU/mice of rVV Nluc/Scarlet, rVV Nluc/GFP, or WR (**Figure 3**). Then, body weight (**Figure 3A**) and survival (**Figure 3B**) were daily monitored for 14 days. No significant body weight loss was observed in mice inoculated with 10^4^ PFU of rVV Nluc/Scarlet, rVV Nluc/GFP, or WR (**Figure 3A**). However, mice inoculated with 10^5^ PFU of rVV Nluc/Scarlet, rVV Nluc/GFP, or WR, lost 15-20% of their initial body weight (**Figure 3A**). Mice infected with 10^6^ PFU and 10^7^ PFU of rVV Nluc/Scarlet, rVV Nluc/GFP, or WR, drastically lost body weight (**Figure 3A**), and all succumbed to viral infection (**Figure 3B**), with those infected with 10^7^ PFU succumbing faster, compared to the rest of the groups. Notably, the 50% mouse lethal dose (MLD_50_) of rVV Nluc/Scarlet and rVV Nluc/GFP (10^5.25^ PFU) was comparable to the MLD_50_ of WR (10^5^ PFU) (**Figure 3B**). These findings indicate that insertion of reporter genes Nluc/Scarlet or Nluc/GFP did not affect the virulence of rVV Nluc/Scarlet and rVV Nluc/GFP, respectively, as compared to WR.

**Figure 3.**
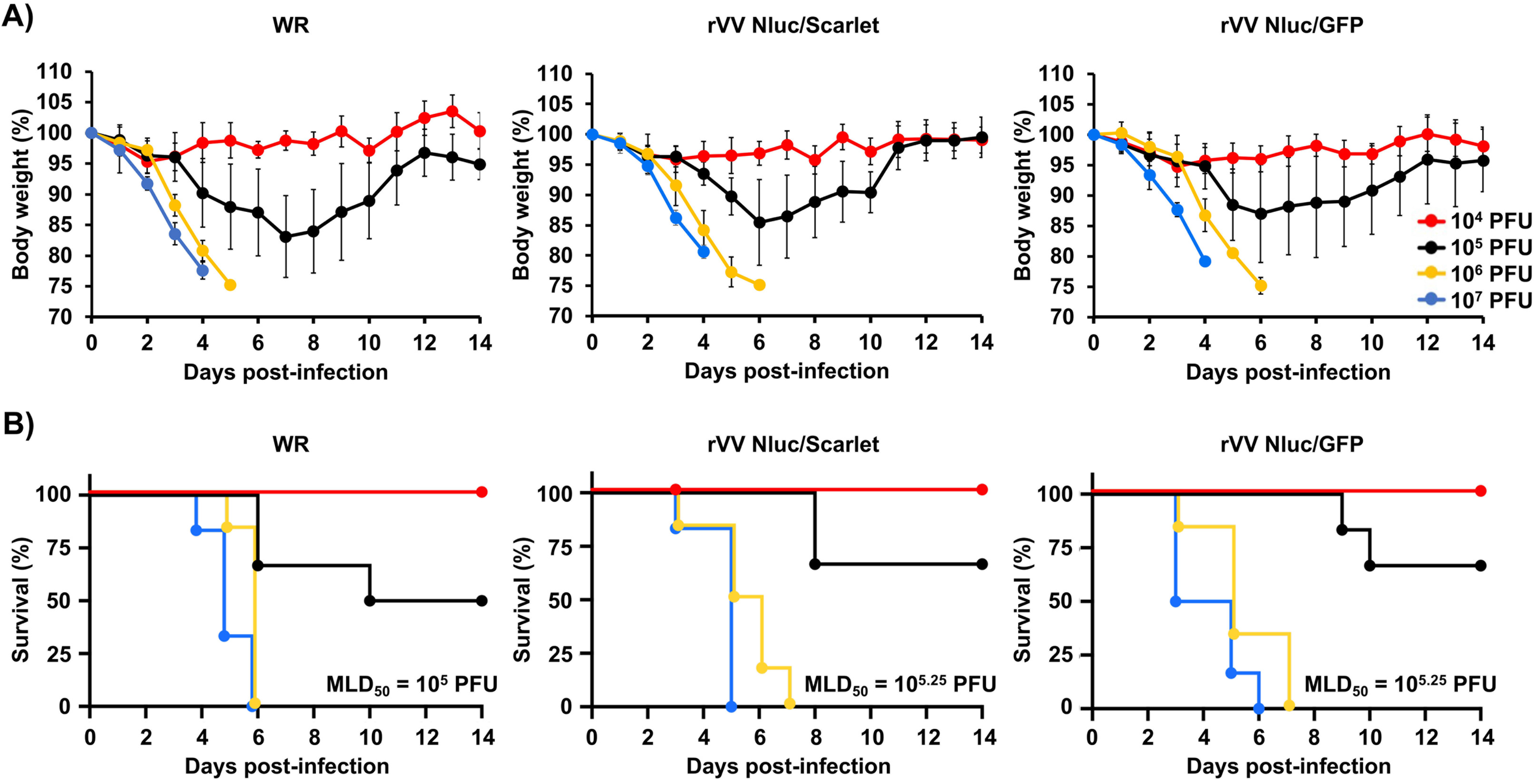
Virulence of bi-reporter rVV in mice: Six-to-eight-week-old female BALB/c mice (n = 6) were i.n. infected with the indicated viral doses (10^4^, 10^5^, 10^6^, or 10^7^ PFU/mice) of WR (left), rVV Nluc/Scarlet (middle) or rVV Nluc/GFP (right). Body weight loss **(A)** and survival **(B)** were monitored for 14 days. Data represent the mean and standard deviation for each group of mice. The MLD_50_ of each virus is indicated in panel B.

### *In vivo* kinetics of rVV Nluc/Scarlet and rVV Nluc/GFP infection

Since detecting fluorescent proteins *in vivo* using *in vivo* imaging is challenging because its intensity often quenches in live tissues (21, 41), we focused on evaluating the dynamics of rVV Nluc/Scarlet and rVV Nluc/GFP infection in mice by bioluminescence (Nluc). BALB/c mice (n = 4/group) were i.n. infected with 10^4^, 10^5^, 10^6^, and 10^7^ PFU/mice of rVV Nluc/Scarlet, rVV Nluc/GFP, or WR, and bioluminescence was monitored every 2 days, for 14 days (**Figure 4**). Bioluminescent imaging of the same mice infected with 10^4^ (**Figure 4A**), 10^5^ (**Figure 4B**), 10^6^ (**Figure 4C**), and 10^7^ (**Figure 4D**) PFU allowed us to temporally visualize viral infection and determine the bioluminescence signal intensities as total flux (log10 photons per second (p/s) (**Figure 4E**). Nluc expression was readily detected in mice infected with the highest doses (10^6^ and 10^7^ PFU) of rVV Nluc/Scarlet and rVV Nluc/GFP as early as day 2 post-infection, although the Nluc intensity signal was higher in those animals infected with 10^7^ PFU of rVV Nluc/Scarlet and rVV Nluc/GFP. Nluc expression in mice infected with 10^5^ PFU of rVV Nluc/Scarlet and rVV Nluc/GFP was observed after 4 days post-infection, while Nluc signal was detected after 6 days post-infection in mice infected with 10^4^ PFU of rVV Nluc/Scarlet and rVV Nluc/GFP. Peak of Nluc expression in mice infected with 10^7^ PFU of rVV Nluc/Scarlet and rVV Nluc/GFP was observed at 4 days post-infection. In this case, mice could not be monitored at later times post-infection since all of them succumbed to viral infection (**Figures 4D and 4E**). In the case of mice infected with 10^6^ PFU of rVV Nluc/Scarlet and rVV Nluc/GFP, maximum levels of Nluc expression were detected at days 4-6 post-infection and could not be monitored at later times post-infection since all of them succumbed to viral infection (**Figures 4C and 4E**). Mice infected with 10^5^ PFU of rVV Nluc/Scarlet and rVV Nluc/GFP showed maximum levels of Nluc expression between days 4-8 post-infection, and we were able to monitor Nluc levels of expression during the entire experiment since all the mice survived infection (**Figures 4B and 4E**). Finally, mice infected with 10^4^ PFU of rVV Nluc/Scarlet and rVV Nluc/GFP showcased detectable peak in Nluc expression at days 8-12 post-infection (**Figures 4A and 4E**), and all the mice survived viral infection.

**Figure 4.**
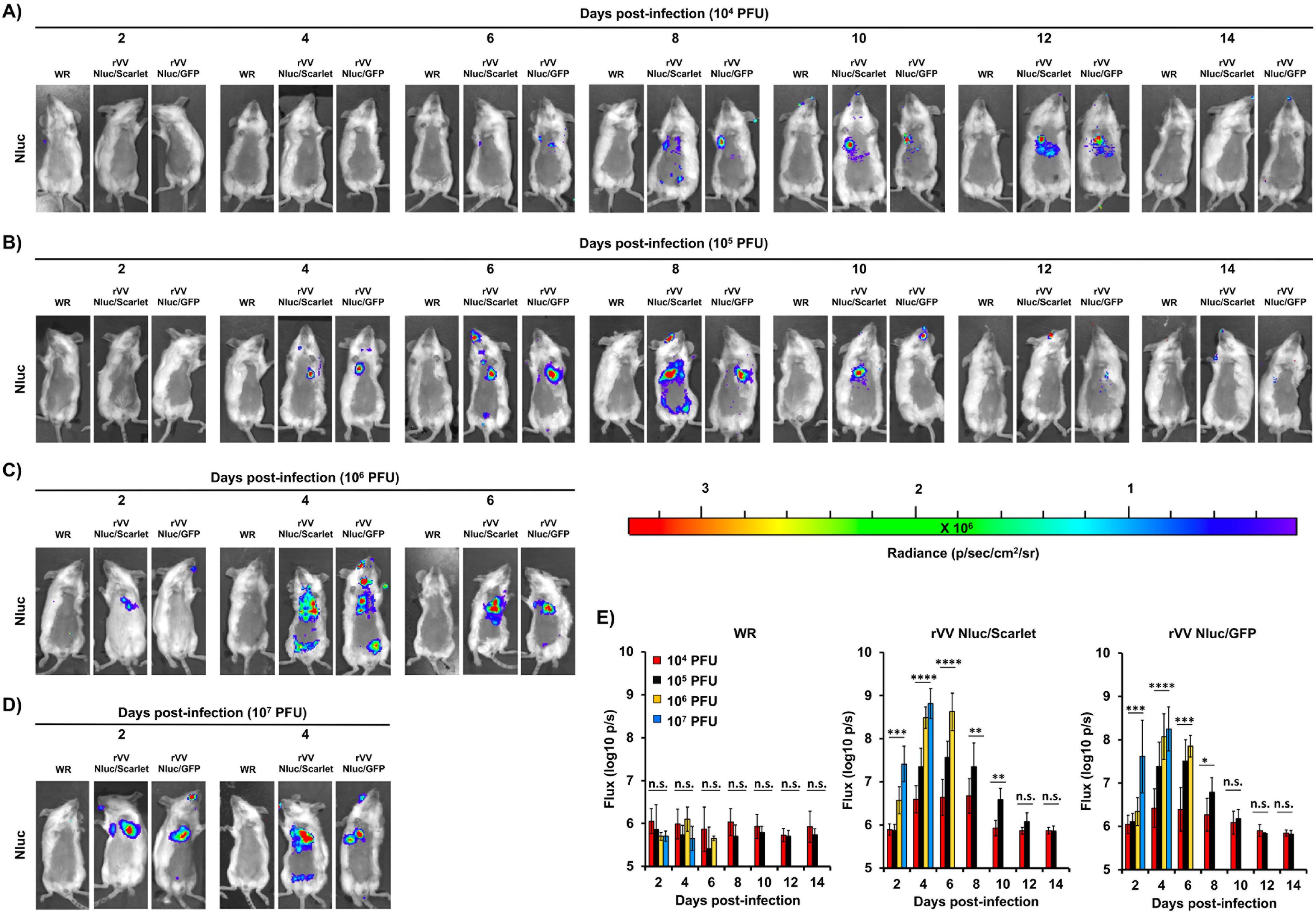
*In vivo* kinetics of bi-reporter expressing rVV: Six-to-eight-week-old female BALB/c mice (n = 4) were infected i.n. with 10^4^ **(A)**, 10^5^ **(B)**, 10^6^ **(C)**, or 10^7^ **(D)** PFU of WR (left), rVV Nluc/Scarlet (middle), or rVV Nluc/GFP (right). Nluc activity was evaluated every 2 days for 14 days (representative images of a single mouse per time point are shown). Radiance (number of photons per second per square centimeter per steradian, p/sec/cm^2^/sr) is shown for each mouse and time point. Bioluminescence signal was quantified and expressed as the total flux (log_10_ protons per second, p/s) **(E).** Error bars indicate the mean and standard deviation of each group of mice. One-way ANOVA was used for statistical analysis; * p < 0.05, ** p < 0.01, *** p < 0.001, **** p < 0.0001 or no significance (n.s.). The line represents the geometric mean.

### *In vivo* fluorescence and Nluc expression, and correlation with viral infection

A major advantage of our dual reporter-expressing rVV is that they harbor both bioluminescent and fluorescent reporter genes. Therefore, either one could be used as a valid surrogate of viral infection. To demonstrate a direct correlation between bioluminescent (Nluc) and fluorescent (Scarlet or GFP) *in vivo* and *ex vivo*, respectively, we infected BALB/c mice (n = 4/group) with 10^4^ (**Figure 5A**), 10^5^ (**Figure 5B**), 10^6^ (**Figure 5C**), and 10^7^ (**Figure 5D**) PFU/mice of rVV Nluc/Scarlet, rVV Nluc/GFP, or WR. We then assessed Nluc expression at days 2, 4, and 6 post-infection using *in vivo* imaging. Immediately after imaging, mice were euthanized, and lungs were collected to quantitate Scarlet (rVV Nluc/Scarlet) or GFP (rVV Nluc/GFP) expression in the lungs using *ex vivo* imaging. Similar to our previous results, we did not detect Nluc expression in mice that were infected with 10^4^ PFU of either rVV Nluc/Scarlet or rVV Nluc/GFP (**Figure 5A**). Some mice infected with 10^4^ PFU of rVV Nluc/Scarlet or rVV Nluc/GFP showed a minimal Nluc signal at day 6 post-infection (**Figure 5A**). Likewise, we could not detect Scarlet or GFP expression in the lungs of mice infected with 10^4^ PFU of rVV Nluc/Scarlet or rVV Nluc/GFP, respectively (**Figure 5A**). Nluc was readily detected in mice inoculated with higher doses of rVV Nluc/Scarlet or rVV Nluc/GFP (10^5^, 10^6^, or 10^7^ PFU). In mice infected with 10^5^ PFU, Nluc was detected by day 4 post-infection (**Figure 5B**). In contrast, Nluc was detected as early as two days post-infection in mice infected with 10^6^ and 10^7^ PFU, respectively (**Figures 5C and 5D**). *Ex vivo* imaging of fluorescent expression in excised lungs correlated with levels of Nluc expression. Scarlet and GFP expression were detected by day 4 post-infection in mice infected with 10^5^ PFU of rVV Nluc/Scarlet or rVV Nluc/GFP (**Figure 5B**); and as early as 2 days post-infection in mice infected with 10^6^ and 10^7^ PFU of rVV Nluc/Scarlet or rVV Nluc/GFP, respectively (**Figures 5C and 5D**). We also determined viral titers from homogenized lungs (**Figure 5E**). As expected, viral titers were higher in mice infected with 10^7^ PFU of rVV Nluc/Scarlet or rVV Nluc/GFP, while the lowest viral titers were observed in mice infected with 10^4^ PFU of rVV Nluc/Scarlet or rVV Nluc/GFP (**Figure 5E**). Median Tissue Culture Infectious Dose (TCID_50_) values in mice infected with 10^4^,10^5^, 10^6^, and 10^7^ PFU of rVV Nluc/Scarlet or rVV Nluc/GFP were within those observed in mice infected with the same viral doses of WR (**Figure 5E**). Lastly, Nluc levels of expression correlated with both Nluc signal detected in whole mice and viral titers (**Figure 5F**).

**Figure 5.**
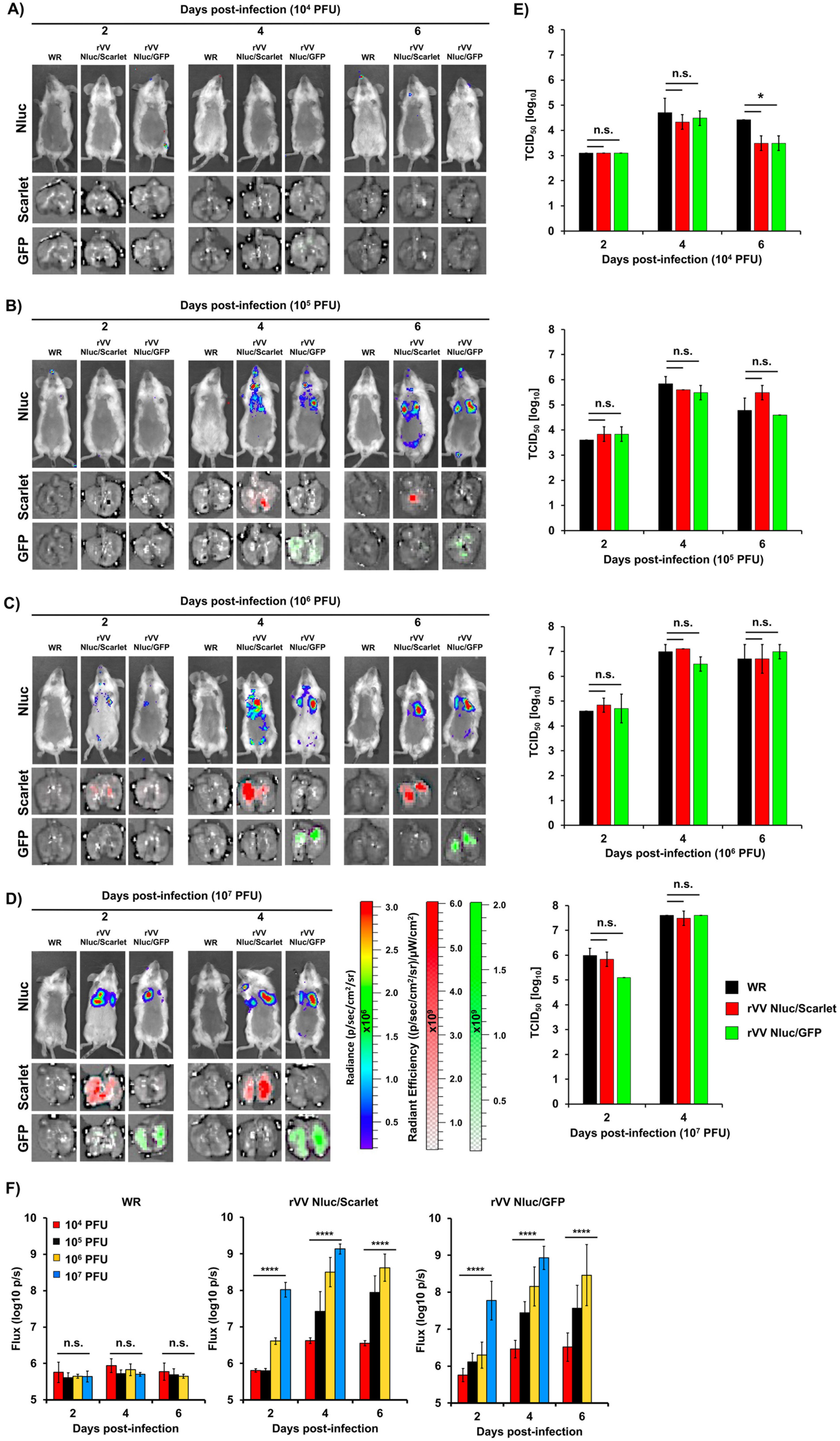
*In vivo* correlation of fluorescence (Scarlet and GFP) and Nluc expression with viral infection: Six-to-eight-week-old female BALB/c mice (n = 4) were i.n. infected with 10^4^ **(A)**, 10^5^ **(B)**, 10^6^ **(C)**, or 10^7^ **(D)** PFU of WR (left), rVV Nluc/Scarlet (middle), or rVV Nluc/GFP (right). Nluc activity (top) was determined at days 2, 4, and 6 post-infection in live mice. Representative images of a single mouse per time point and p/sec/cm^2^/sr values of all mice per time point are shown. After *in vivo* imaging, lungs were harvested to access Scarlet (middle) or GFP (bottom) expression. Viral titers in the lungs of the same mice shown in panels A-D were determined by TCID_50_ (**E**). Error bars indicate the mean and standard deviation of each group of mice. Bioluminescence signal was quantified and expressed as the total flux (log_10_ protons per second, p/s) **(F).** Error bars indicate the mean and standard deviation of each group of mice. The One-way ANOVA was used for statistical analysis; * p < 0.05, ** p < 0.01, *** p < 0.001, **** p < 0.0001 or no significance (n.s.). The line represents the geometric mean.

### Direct visualization of rVV Nluc/Scarlet and rVV Nluc/GFP in mice lungs

One of the practical applications of viruses expressing fluorescent reporter genes is the rapid and easy detection of the pathogen without the need for additional reagents to identify the presence of the virus in infected cells. In addition, the resolution of microscopical imaging using fluorescent reporters allows identifying individual positive cells without the need for antibody staining. Thus, we decided to visualize the spatial distribution of the VV and their cellular targets in formalin-fixed, paraffin-embedded lung sections of BALB/c mice (n = 4) infected with 10^7^ PFU of rVV Nluc/Scarlet or rVV Nluc/GFP at days 2 and 4 post-infection by examining Scarlet and GFP expression, respectively (**Figure 6**). As expected, we did not find a fluorescent signal in the lungs of mock-infected mice or in mice infected with WR (**Figure 6A**). However, lung sections from rVV Nluc/Scarlet- and rVV Nluc/GFP-infected mice displayed red and green fluorescent signals, respectively. Although some epithelial bronchi were virus-free, we observed a progressive increase in fluorescent signals in the airways between days 2 and 4 post-infection (**Figure 6A**). As a result of the fixation process, in certain instances, fluorescence intensity might be drastically diminished. To overcome this problem, researchers have used antibodies specific for fluorescent proteins to identify sites of viral infection. Therefore, we stained lung sections of mice infected with rVV Nluc/Scarlet or rVV Nluc/GFP with antibodies specific for Scarlet (mCherry) or GFP, respectively (**Figure 6B**). Of note, lung samples from mock- or WR-infected mice were negative for Scarlet and GFP (**Figure 6B**). In contrast, Scarlet and GFP expression were enhanced by specific antibodies and were mainly located in the airways of the lung tissues (**Figure 6B**). As we predicted, viral infection was detected using a specific antibody against VV in the lung sections of all mice groups infected with WR, rVV Nluc/Scarlet, or rVV Nluc/GFP, but not in mock-infected animals (**Figure 6C**). The staining patterns observed after direct visualization of Scarlet or GFP expression from reporter-expressing viruses (**Figure 6A**), incubation with antibodies against fluorescent proteins (**Figure 6B**), or VV (**Figure 6C**) were similar and indicative of epithelial cell infections in the bronchial airways (**Figures 6A-6C**). Lastly, a morphometric analysis of the area infected by WR, rVV Nluc/Scarlet, or rVV Nluc/GFP at days 2 and 4 post-infection revealed a significant 4-fold increase in the percentage of area covered by the fluorescent signal on day 4 post-infection (**Figure 6D**).

**Figure 6.**
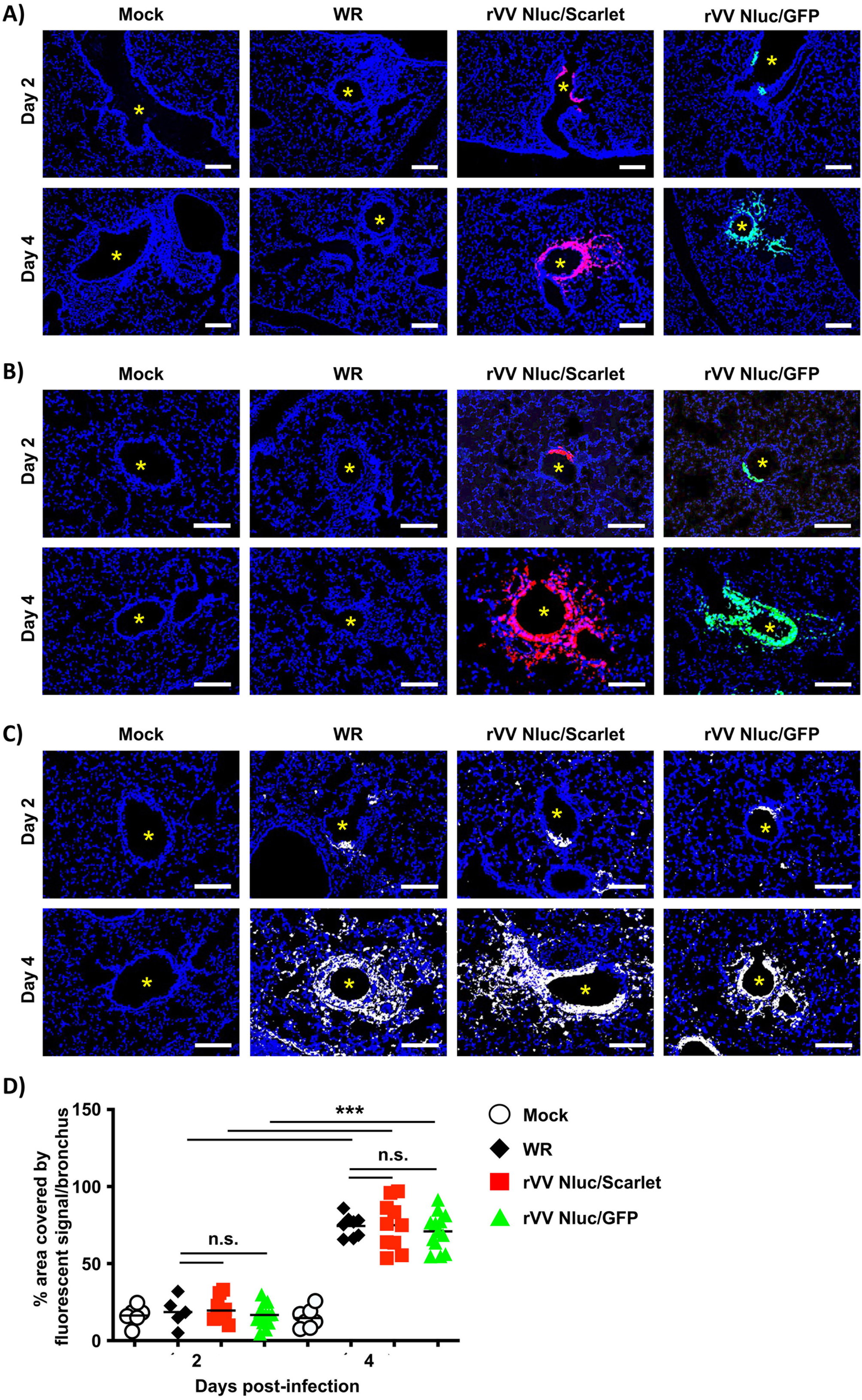
Direct visualization of viral infection in lungs of BALB/c mice infected with bi-reporter expressing rVV: Six-to-eight-weeks-old female BALB/c mice (n = 4) were mock-infected or infected with 10^7^ PFU of WR, rVV Nluc/Scarlet, or rVV Nluc/GFP. Mice were sacrificed on days 2 and 4 post-infection and lungs were harvested to assess fluorescent Scarlet and GFP expression. Representative images from lungs of mock-infected and mice infected with WR, rVV Nluc/GFP, or rVV Nluc/Scarlet viruses on days 2 and 4 without immunostaining **(A),** stained with antibodies against Scarlet or GFP **(B)** or a rabbit polyclonal antibody against VV **(C)** are shown. Airways are noted with an asterisk. Scale bars 100 μm. Morphometric analyses of the infected bronchi in these experiments are also shown **(D)**. One-way ANOVA was used for statistical analysis; * p < 0.05, ** p < 0.01, *** p < 0.001, or no significance (n.s.). The line represents the geometric mean.

### Pathological changes in mice lungs infected with dual reporter-expressing rVV

We next evaluated the ability of rVV Nluc/Scarlet and rVV Nluc/GFP to induce pathological changes in the lungs of infected (10^7^ PFU) BALB/c mice (n = 4) and compared to those caused by WR infection, and mock-infected animals (**Figure 7**). Lungs of mice infected with either rVV Nluc/Scarlet, rVV Nluc/GFP, or WR, showed mild to moderate bronchopneumonia (**Figure 7A**). Blind assessment of pathological pulmonary features confirmed our preliminary observations and showed comparable inflammatory cell infiltration around bronchi and blood vessels (**Figures 7A and 7B**, blue arrows). Accumulation of inflammatory cells in the intra-alveolar space was significantly higher in WR-infected mice compared to rVV Nluc/Scarlet and rVV Nluc/GFP infected animals at day 2 post-infection, but was not statistically different by day 4 post-infection (**Figures 7A and 7C**, green arrowheads). Bronchial epithelial cell necroses were similar in mice infected with WR and dual reporter expressing rVV Nluc/Scarlet, or rVV Nluc/GFP (**Figures 7A and 7D**, black arrows).

**Figure 7.**
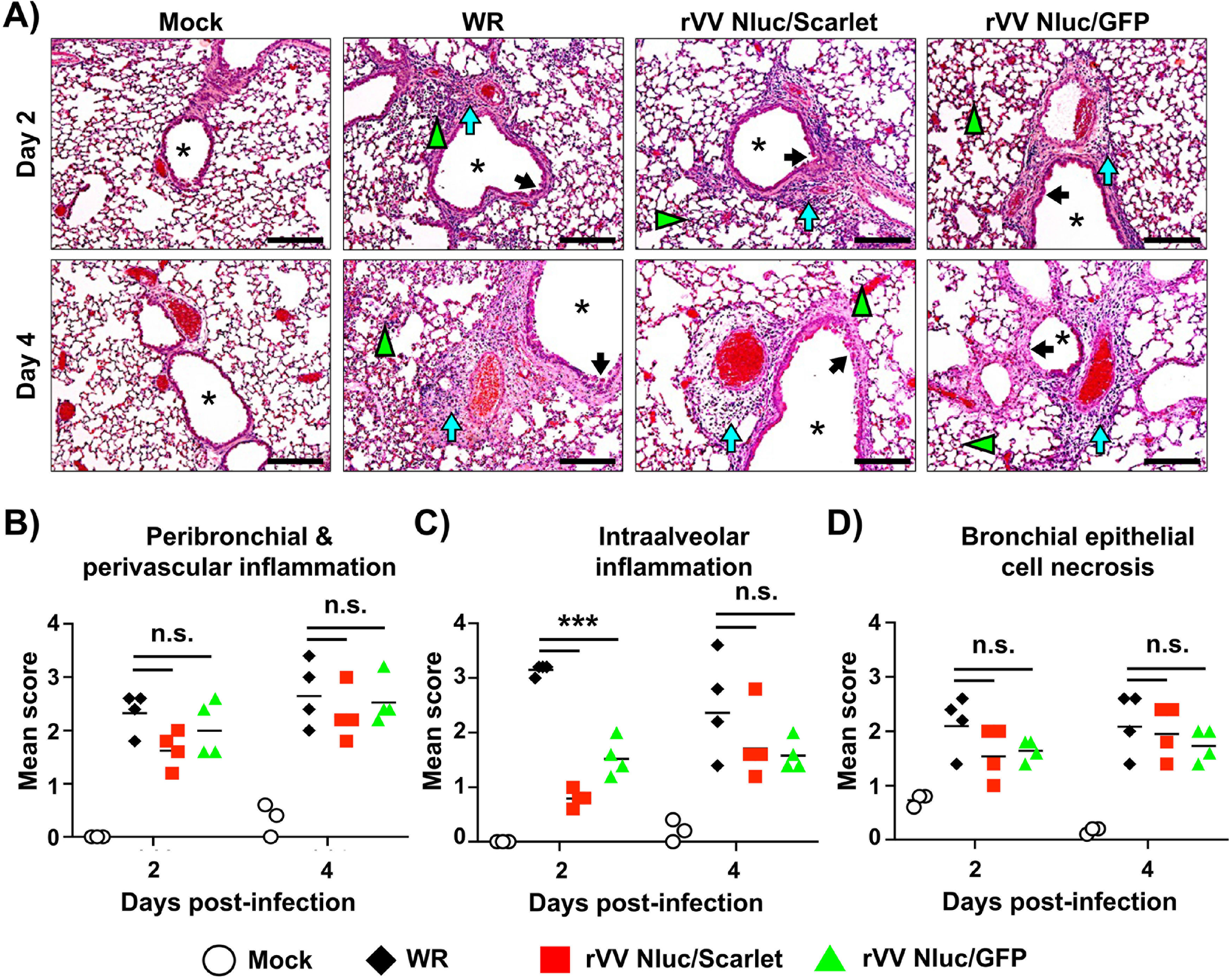
Pathological changes in the lungs of mice infected with the bi-reporter expressing rVV: Six-to-eight-weeks–old female BALB/c mice (n = 4) were mock-infected or infected i.n. with 10^7^ PFU of WR, rVV Nluc/Scarlet, or rVV Nluc/GFP. At days 2 and 4 post-infection, mice were sacrificed, and lung samples were collected for blinded histopathologic examination **(A)** and scoring **(B-D)**. Lung infected with WR, rVV Nluc/Scarlet, and rVV Nluc/GFP showed multifocal to locally extensive mild to moderate peribronchial & perivascular inflammation **(A,** green arrow), intra-alveolar inflammations **(A,** blue arrowhead), and bronchial epithelial cell necrosis **(A,** black arrow). Mock-infected mice showed no lesions. Airways are noted with an asterisk. Scale bars 50 μm. The histopathologic lesion scoring from peribronchial & perivascular inflammations **(B),** intra-alveolar inflammation **(C)** and bronchial epithelial cell necrosis **(D)** were graded base on lesion severity as follows: grade 0 = no histopathological lesions observed; grade 1 = mild; grade 2 = moderate; grade 3 = marked; and grade 4 = severe. One-way ANOVA was used for statistical analysis; * p < 0.05, ** p < 0.01, *** p < 0.001, or no significance (n.s.). The line represents the geometric mean.

### Identification of the cell type targeted by VV in BALB/c mice

To identify the cellular targets of VV in infected mice, we enumerated and calculated the proportion of immune cell subsets (CD45^-^) and stromal cells (CD45^+^) in the lung of mice infected with 10^7^ PFU of rVV Nluc/GFP by flow cytometry (**Figure 8**). We excluded dead cells and doublets from the flow cytometry analysis to detect exclusively live macrophages (CD45^+^CD11b^+^CD64^+^), neutrophils (CD45^+^CD11b^+^Ly6G^+^), monocyte-derived dendritic cells (CD45^+^CD11c^+^I-A^b+^CD11b^+^Ly6C^int^), monocyte undifferentiated macrophages (CD45^+^CD11c^+^IA^b-/low^CD11b^+^Ly6C^int^) and interstitial macrophages (CD45^+^CD11b^+^IA^b+^ CD6^int/hi^) that have active viral infection by assessing GFP reporter expression (**Supplementary Figure 1**). Consistent with the progressive spread of viral infection in the airways, we found a significant increase in the proportion of CD45^-^ stromal cells infected from days 2 to 4 post-infection (**Figure 8A**, day 2; 2.7% CD45^-^ GFP^+^ vs. day 4; 52% CD45^-^GFP^+^). The frequency and total number of monocytes undifferentiated macrophages (day 2; 40.5% vs. day 4; 31.3%), monocyte-derived dendritic cells (day 2; 31.5% vs. day 4; 11.3%) and interstitial macrophages (day 2; 12.3% vs. day 4; 3.4%) infected by rVV Nluc/GFP was lower at day 4 compared to day 2 post-infection (**Figures 8A and 8B**). In contrast, infected macrophages (day 2; 8% vs. day 4; 29.2%) and neutrophils (day 2; 7.5% vs. day 4; 24.8%) increased at day 4 post-infection, which may be associated with the differentiation of monocytes into macrophages in the infected lungs (**Figures 8A and 8B**). Yet, both rVV Nluc/GFP and WR efficiently induced an innate immune response in the lung of infected mice. rVV Nluc/GFP induced significant accumulation of macrophages and interstitial macrophages (day 2), monocyte-derived dendritic cells, and monocytes undifferentiated macrophages (day 4) compared to WR-infected mice. Intriguingly, we observed a decrease of total macrophages and neutrophils at day 4 post-infection that may be associated with the increase in viral infection (**Figure 8B**).

**Figure 8.**
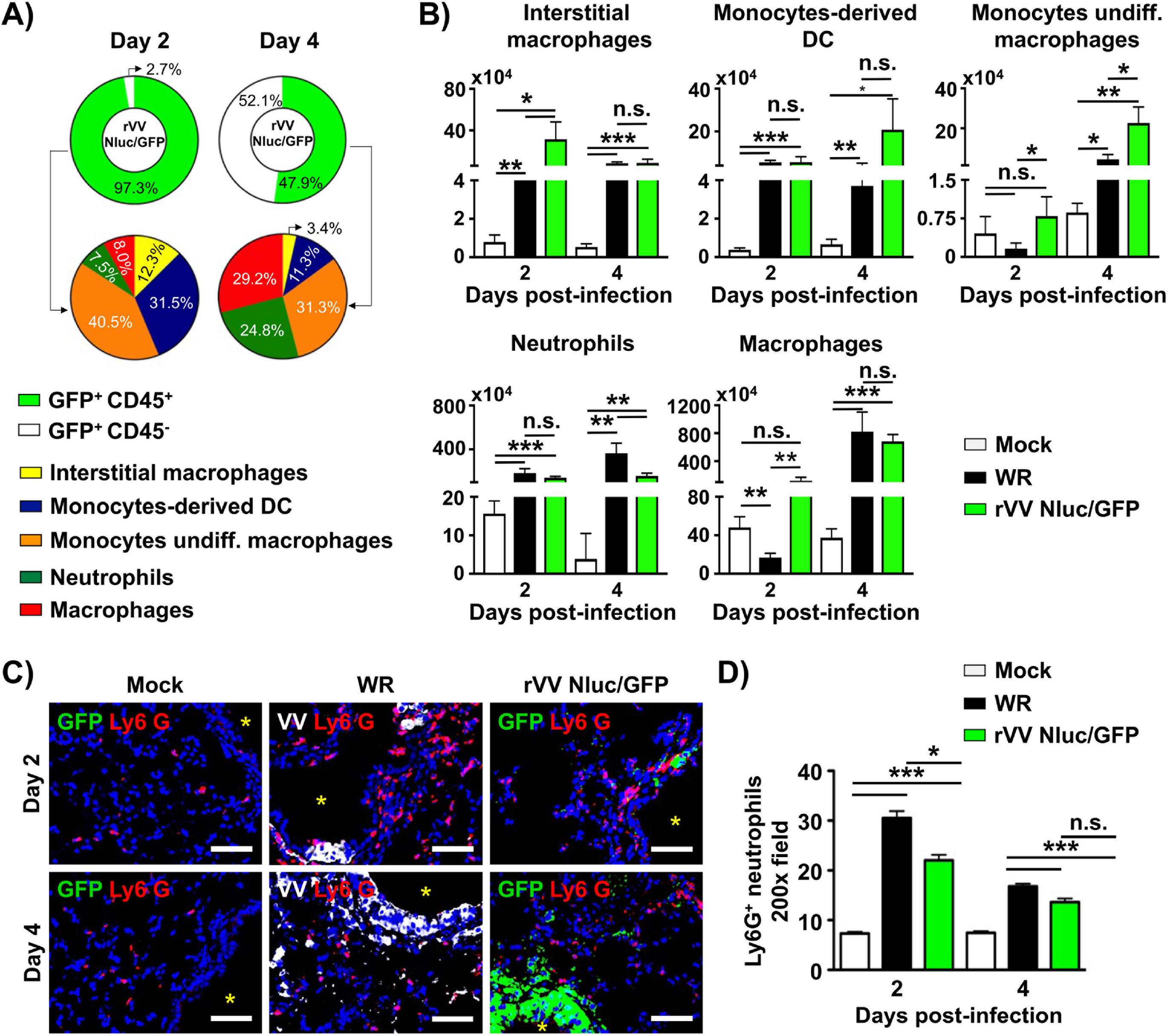
Identification of cells infected with rVV: Six-to-eight-weeks-old female BALB/c mice (n = 4) were mock-infected or infected i.n. with 10^7^ PFU of WR or rVV Nluc/GFP. At days 2 and 4 post-infection, lungs were harvested, enzymatically and mechanically dissociated into single-cell suspensions, and the composition of GFP-positive cells was evaluated by flow cytometry. Pie charts represent distinguished CD45^-^ stromal cells, CD45^+^ hematopoietic cells, and innate immune cells responsive to rVV Nluc/GFP from infected mice at days 2 or 4 post-infection **(A).** The total number of interstitial macrophages, monocyte-derived DC, monocytes/undifferentiated macrophages, neutrophils and macrophages in the lungs from mock-infected and infected with WR or rVV Nluc/GFP mice is shown (**B).** The spatial distribution of infiltrating neutrophils in mock-infected and WR or rVV Nluc/GFP mice are shown **(C)**. Ly6G+ neutrophils are shown in red. WR infected cells are shown in white. rVV Nluc/GFP infected cells are shown in green. Scale bars 100 µm. Airways are noted with an asterisk. Representative images are shown. The morphometric analysis depicts the average number of Ly6G+ neutrophils per 200x fields in the site of infection **(D).** The One-way ANOVA was used for statistical analysis; * p < 0.05, ** p < 0.01, *** p < 0.001, or no significance (n.s.).

Next, we performed immunofluorescence staining to identify the spatial distribution of innate cells at the site of infection. To visualize the location of neutrophils relative to the infected airways, we stained lung sections with antibodies specific for Ly6G, VV, and GFP to detect neutrophils, WR, and rVV Nluc/GFP, respectively. In the lungs of mock-infected mice, we observed most neutrophils accumulated in the interstitial space, while very few were near the airways (**Figure 8C**, left panels). In contrast, many neutrophils accumulated around airways in mice infected with WR or rVV Nluc/GFP (**Figure 8C**, middle and right panels). Consistent with our preliminary observations, the number of neutrophils significantly increased around the airways in mice infected with WR or rVV Nluc/GFP (**Figure 8D**). The number of neutrophils infiltrating the peri-bronchial areas was similar in mice infected with either WR or rVV Nluc/GFP (**Figure 8D**).

### Stability of rVV Nluc/Scarlet and rVV Nluc/GFP

A critical concern with reporter-expressing recombinant viruses is their genetic instability that might lead to the loss of correct marker gene expression. To test the stability of our bi-reporter constructs, we serially passaged both, rVV Nluc/Scarlet and rVV Nluc/GFP, as well as WR, in cultured cells for a total of 5 passages. Next, virus progeny obtained from passage 5 was evaluated by plaque assay and monitored for reporter gene expression (**Figure 9**). Results indicate that rVV Nluc/Scarlet and rVV Nluc/GFP were genetically stable since both recombinant viruses displayed the expected fluorescent (Scarlet and GFP) and bioluminescence (Nluc) expression (**Figure 9**).

**Figure 9.**
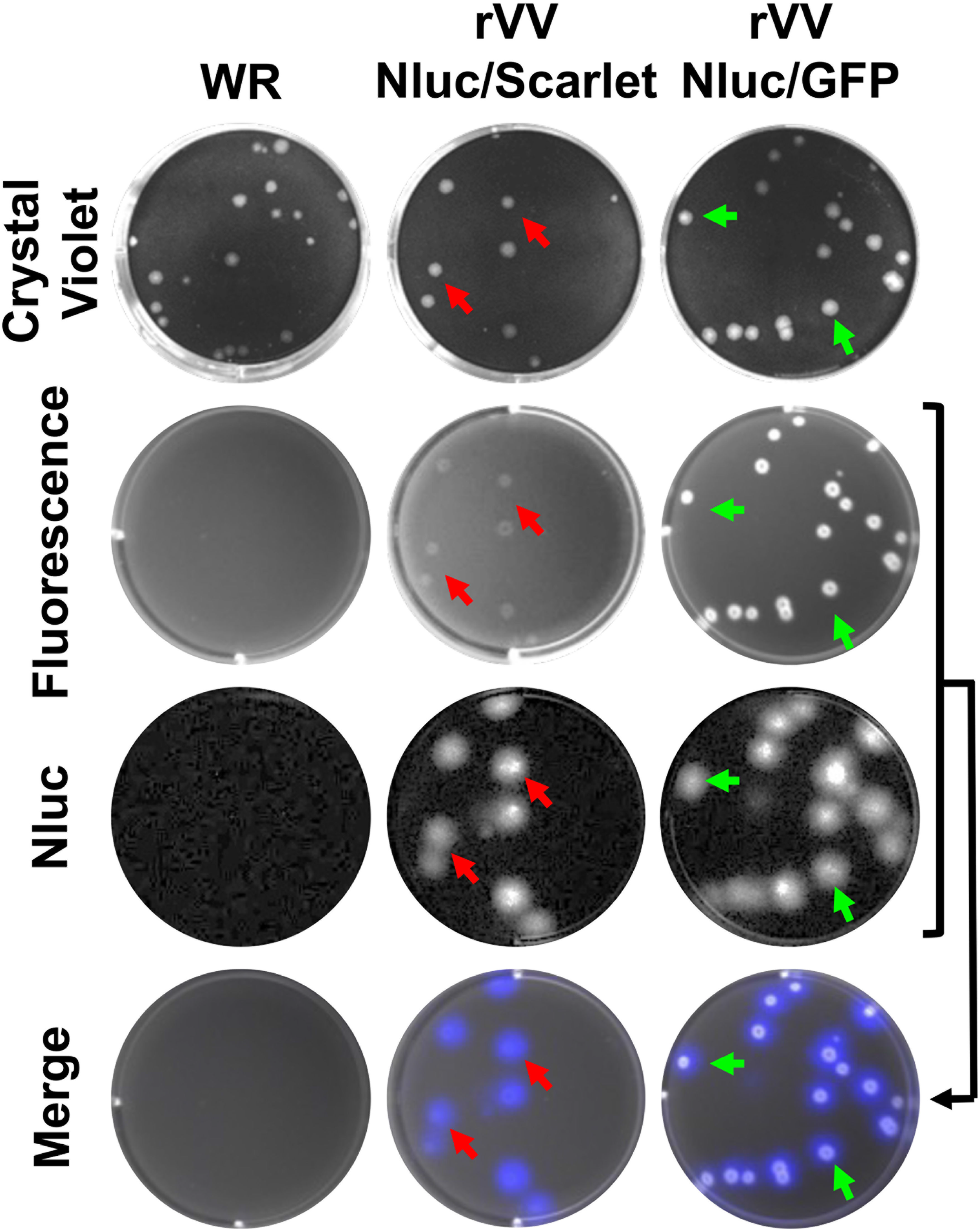
Stability of rVV Nluc/Scarlet and rVV Nluc/GFP: rVV Nluc/Scarlet and rVV Nluc/GFP were serially passaged 5 times in BSC-1 cells. Parental WR was also passaged 5 times as internal control. Passage 5 WR, rVV Nluc/Scarlet and rVV Nluc/GFP were analyzed by plaque assay in BSC-1. Representative images of WR (left), rVV Nluc/Scarlet (middle) and rVV Nluc/GFP (right) plaques in BSC-1 cells at 2 days post-infection are shown. Plaques were detected by crystal violet staining, Scarlet or GFP (fluorescence), and luciferase (Nluc) expression. Merged images are shown at the bottom. Red (rVV Nluc/Scarlet) and green (rVV Nluc/GFP) arrows show the co-localization of Scarlet or GFP fluorescence, respectively, with crystal violet and Nluc viral plaques.

## DISCUSSION

Previous studies, including ours, have described reporter-expressing rVV encoding either fluorescent or luciferase reporter genes to study the biology of VV and to evaluate the efficacy of new antivirals or NAbs to inhibit or neutralize, respectively, viral infection (24, 42, 43). However, and based on the different properties of fluorescent and luciferase proteins, the use of these reporter-expressing rVV is limited to the properties of the reporter gene (15, 41, 44). *In vitro*, fluorescent proteins represent a better option to detect the presence of rVV in infected cells (14–17). However, luciferase proteins represent a better tool for quantitative purposes (19–21, 45). *In vivo*, luciferase reporter genes represent the best option for whole-animal imaging (20, 41), while fluorescent proteins are better for *ex vivo* imaging and identifying viral cellular targets (15, 46). Thus, rVV expressing only one reporter gene does not allow to take advantage of the different properties of fluorescent and luciferase proteins and, therefore, the properties of the reporter fluorescent or luciferase proteins must be carefully considered in the experimental design.

Here, we have described the generation of novel rVV stably expressing two different reporter fluorescent and luciferase genes to overcome the current limitations of using single reporter-expressing rVV (**Figure 1**). We previously demonstrated the feasibility and advantages of generating bi-reporter influenza A virus (IAV) (37, 38) and the prototype arenavirus lymphocytic choriomeningitis virus (LCMV) (47). This new dual reporter-expressing rVV possess the advantages of both fluorescent and luciferase reporters, which could provide promising applications for the study of VV *in vitro* and *in vivo*. Notably, the expression of two foreign genes from different loci in the viral genome opens the possibility of improving the vaccine vector capability features of VV.

We found that our bi-reporter expressing rVV have similar growth kinetics and plaque phenotype in cultured cells than the parental WR strain (**Figure 2**). Importantly, only rVV expressing both reporter genes were detectable by fluorescence microscopy or bioluminescence due to its expression of GFP or Scarlet, and Nluc, respectively. Infection with rVV expressing both reporter genes was visualized in real time, without the need of complex protocols (e.g. staining with antibodies) needed to detect the presence of parental VV strains in infected cells. Moreover, reporter gene expression correlated to viral replication, demonstrating the feasibility of using either the fluorescent or luciferase reporter genes as valid surrogates of infection. In 2018, the United States FDA-approved the use of tecovirimat (TPOXX) to treat poxvirus infections (3). However, additional antivirals for the efficient treatment of poxvirus infections are urgently needed. Thus, discovering novel antivirals and implementing rapid and sensitive screening assays amenable to HTS will accelerate the identification and characterization of novel anti-poxvirus compounds. Antiviral drug discovery against poxvirus infection will benefit of rVV expressing both fluorescent and luciferase proteins where identification of antivirals will be based on orthogonal assays based on inhibition of both reporter genes, similar to our previous studies with bi-reporter IAV (38) and LCMV (47). Likewise, these bi-reporter expressing rVV represent an excellent option to assess the neutralizing capacity of antibodies in HTS settings, as previously described for IAV (38, 48).

*In vivo*, the dual reporter-expressing rVV retained parental WR-like morbidity and mortality, with animals losing body weight and succumbing to viral infection, respectively, similar to those infected with WR (**Figure 3**). Notably, the bi-reporter expressing rVV represent a powerful tool to visualize viral infection *in vivo* (Nluc) (**Figure 4**) or *ex vivo* (Scarlet and GFP) (**Figure 5**). When we imaged mice infected with rVV Nluc/Scarlet or rVV Nluc/GFP using IVIS, Nluc signal was easily detectable and quantifiable, revealing the dynamics of VV infection using different doses of viral inoculum (**Figure 4**). Importantly, Nluc activity was time and viral dose dependent and allowed us to longitudinally follow and measure viral replications dynamics in the same live animals during the progression of natural viral infection. Likewise, the excised lungs from mice infected with rVV Nluc/Scarlet and rVV Nluc/GFP displayed Scarlet or GFP expression, respectively, where fluorescent signals were also time and viral dose-dependent (**Figure 5**). In addition, fluorescent protein expression allowed us to identify individually infected cells using fluorescent imaging and flow cytometry (**Figure 6**) and to assess the pathology of viral infections in the lungs of infected mice (**Figure 7**). Importantly, we observed a spatial and temporal correlation between Nluc and fluorescent expression (Scarlet or GFP), which correlated with viral replication in the lungs of infected mice (**Figure 5**). Finally, we identified the major primary cellular targets of VV *in vivo* by combining the identification of GFP+ cells with flow cytometry-based approaches (**Figure 8**).

Altogether, rVV stably expressing both fluorescent and luciferase reporter genes represent an excellent tool for studying the biology of VV *in vitro* or *in vivo*. Expression of both reporter genes adds the advantages of using individual fluorescent- or luciferase-expressing rVV and represent an excellent option for the identification of therapeutics against poxviruses. Moreover, our studies with VV provide proof-of-concept for the feasibility of using a similar approach with other poxviruses, or DNA viruses, to facilitate their study in cultured cells or in validated animal models of infection.

## ACKNOWLEDGMENTS

We want to thank BEI Resources for providing reagents. We also want to thank Drs. Hiller and Alcamí for providing antibodies. This work was supported by grants E-RTA2014-00006, RTA2017-0066 from Ministerio de Economía y Competitividad and Ministerio de Ciencia, Innovación y Universidades as part of the Plan Estatal de Investigación Científica, Desarrollo e Innovación Tecnológica, and grant COV20-00901 from Instituto de Salud Carlos III (ISCIII).

## SUPPLEMENTARY FIGURE LEGENDS

**Supplementary Figure 1. Flow cytometry gating strategy used to identify different cell subsets during rVV Nluc/GFP infection:** Six-to-eight-weeks-old female BALB/c mice (n = 4) were i.n. infected with 10^7^ PFU of rVV Nluc/GFP. Lungs were harvested on days 2 and 4 post-infection, enzymatically digested, and mechanically disrupted to obtain single cell suspensions. After gating in live single GFP^+^ cells a stepwise gating strategy was used to distinguish subsets of virus-infected cells based on CD45^+^ expression and staining with specific antibodies to identify and enumerate macrophages (CD64^+^ CD11b^+^), neutrophils (CD64^-^ CD11b^+^ Ly6G^+^), monocyte-derived DC (CD11b^+^1-A^b+^Ly6C^+^), monocytes/undifferentiated macrophages (CD11b^+^CD64^+^I-A^b-/low^Ly6C^+/-^), and interstitial macrophages (CD11b^+^1-A^b+^ CD64^int/hi^). Representative plots for each immune subset are shown (n = 4 mice/group).

## REFERENCES

1. Moss B. 2013. Poxvirus DNA replication. Cold Spring Harb Perspect Biol 5.

2. Shailubhai K. 2003. Bioterrorism: a new frontier for drug discovery and development. IDrugs 6:773–80.

3. Merchlinsky M, Albright A, Olson V, Schiltz H, Merkeley T, Hughes C, Petersen B, Challberg M. 2019. The development and approval of tecoviromat (TPOXX. Antiviral Res 168:168–174.

4. Delaune D, Iseni F. 2020. Drug Development against Smallpox: Present and Future. Antimicrob Agents Chemother 64.

5. Sklenovská N, Van Ranst M. 2018. Emergence of Monkeypox as the Most Important Orthopoxvirus Infection in Humans. Front Public Health 6:241.

6. Smith GL, Benfield CTO, Maluquer de Motes C, Mazzon M, Ember SWJ, Ferguson BJ, Sumner RP. 2013. Vaccinia virus immune evasion: mechanisms, virulence and immunogenicity. J Gen Virol 94:2367–2392.

7. Moss B. 2016. Membrane fusion during poxvirus entry. Semin Cell Dev Biol 60:89–96.

8. Moss B. 2012. Poxvirus cell entry: how many proteins does it take? Viruses 4:688–707.

9. Johnson MC, Damon IK, Karem KL. 2008. A rapid, high-throughput vaccinia virus neutralization assay for testing smallpox vaccine efficacy based on detection of green fluorescent protein. J Virol Methods 150:14–20.

10. Rozelle DK, Filone CM, Dower K, Connor JH. 2014. Vaccinia reporter viruses for quantifying viral function at all stages of gene expression. J Vis Exp doi:10.3791/51522.

11. Dower K, Rubins KH, Hensley LE, Connor JH. 2011. Development of Vaccinia reporter viruses for rapid, high content analysis of viral function at all stages of gene expression. Antiviral Res 91:72–80.

12. Bartee E, Mohamed MR, Lopez MC, Baker HV, McFadden G. 2009. The addition of tumor necrosis factor plus beta interferon induces a novel synergistic antiviral state against poxviruses in primary human fibroblasts. J Virol 83:498–511.

13. Ward BM, Moss B. 2001. Visualization of intracellular movement of vaccinia virus virions containing a green fluorescent protein-B5R membrane protein chimera. J Virol 75:4802–13.

14. Breen M, Nogales A, Baker SF, Perez DR, Martinez-Sobrido L. 2016. Replication-Competent Influenza A and B Viruses Expressing a Fluorescent Dynamic Timer Protein for In Vitro and In Vivo Studies. PLoS One 11:e0147723.

15. Breen M, Nogales A, Baker SF, Martínez-Sobrido L. 2016. Replication-Competent Influenza A Viruses Expressing Reporter Genes. Viruses 8.

16. Nogales A, Baker SF, Martínez-Sobrido L. 2015. Replication-competent influenza A viruses expressing a red fluorescent protein. Virology 476:206–16.

17. Nogales A, Rodriguez-Sanchez I, Monte K, Lenschow DJ, Perez DR, Martinez-Sobrido L. 2015. Replication-competent fluorescent-expressing influenza B virus. Virus Research 213:69–81.

18. Stacer AC, Nyati S, Moudgil P, Iyengar R, Luker KE, Rehemtulla A, Luker GD. 2013. NanoLuc reporter for dual luciferase imaging in living animals. Mol Imaging 12:1–13.

19. Tran V, Poole DS, Jeffery JJ, Sheahan TP, Creech D, Yevtodiyenko A, Peat AJ, Francis KP, You S, Mehle A. 2015. Multi-Modal Imaging with a Toolbox of Influenza A Reporter Viruses. Viruses 7:5319–27.

20. Czako R, Vogel L, Lamirande EW, Bock KW, Moore IN, Ellebedy AH, Ahmed R, Mehle A, Subbarao K. 2017. In Vivo Imaging of Influenza Virus Infection in Immunized Mice. MBio 8.

21. Heaton NS, Leyva-Grado VH, Tan GS, Eggink D, Hai R, Palese P. 2013. In vivo bioluminescent imaging of influenza a virus infection and characterization of novel cross-protective monoclonal antibodies. J Virol 87:8272–81.

22. Kelkar M, De A. 2012. Bioluminescence based in vivo screening technologies. Curr Opin Pharmacol 12:592–600.

23. Hall MP, Unch J, Binkowski BF, Valley MP, Butler BL, Wood MG, Otto P, Zimmerman K, Vidugiris G, Machleidt T, Robers MB, Benink HA, Eggers CT, Slater MR, Meisenheimer PL, Klaubert DH, Fan F, Encell LP, Wood KV. 2012. Engineered luciferase reporter from a deep sea shrimp utilizing a novel imidazopyrazinone substrate. ACS Chem Biol 7:1848–57.

24. Dominguez J, Lorenzo MM, Blasco R. 1998. Green fluorescent protein expressed by a recombinant vaccinia virus permits early detection of infected cells by flow cytometry. J Immunol Methods 220:115–21.

25. Sanchez-Puig JM, Sanchez L, Roy G, Blasco R. 2004. Susceptibility of different leukocyte cell types to Vaccinia virus infection. Virol J 1:10.

26. Lam AJ, St-Pierre F, Gong Y, Marshall JD, Cranfill PJ, Baird MA, McKeown MR, Wiedenmann J, Davidson MW, Schnitzer MJ, Tsien RY, Lin MZ. 2012. Improving FRET dynamic range with bright green and red fluorescent proteins. Nat Methods 9:1005–12.

27. Lorenzo MM, Sánchez-Puig JM, Blasco R. 2019. Genes A27L and F13L as Genetic Markers for the Isolation of Recombinant Vaccinia Virus. Sci Rep 9:15684.

28. Bindels DS, Haarbosch L, van Weeren L, Postma M, Wiese KE, Mastop M, Aumonier S, Gotthard G, Royant A, Hink MA, Gadella TW. 2017. mScarlet: a bright monomeric red fluorescent protein for cellular imaging. Nat Methods 14:53–56.

29. Lorenzo MM, Galindo I, Blasco R. 2004. Construction and isolation of recombinant vaccinia virus using genetic markers. Methods Mol Biol 269:15–30.

30. Chakrabarti S, Sisler JR, Moss B. 1997. Compact, synthetic, vaccinia virus early/late promoter for protein expression. Biotechniques 23:1094–7.

31. Cotter CA, Earl PL, Wyatt LS, Moss B. 2015. Preparation of Cell Cultures and Vaccinia Virus Stocks. Curr Protoc Microbiol 39:14A.3.1–14A.3.18.

32. National Research Council (U.S.). Committee for the Update of the Guide for the Care and Use of Laboratory Animals., Institute for Laboratory Animal Research (U.S.), National Academies Press (U.S.). 2011. Guide for the care and use of laboratory animals, 8th ed. National Academies Press, Washington, D.C.

33. Rodriguez L, Nogales A, Martínez-Sobrido L. 2017. Influenza A Virus Studies in a Mouse Model of Infection. J Vis Exp doi:10.3791/55898.

34. Nogales A, Martinez-Sobrido L, Chiem K, Topham DJ, DeDiego ML. 2018. Functional Evolution of the 2009 Pandemic H1N1 Influenza Virus NS1 and PA in Humans. Journal of Virology 92.

35. Nogales A, Baker SF, Ortiz-Riano E, Dewhurst S, Topham DJ, Martinez-Sobrido L. 2014. Influenza A Virus Attenuation by Codon Deoptimization of the NS Gene for Vaccine Development. Journal of Virology 88:10525–40.

36. Reed LJ, Muench H. 1938. A SIMPLE METHOD OF ESTIMATING FIFTY PER CENT ENDPOINTS12. American Journal of Epidemiology 27:493–497.

37. Chiem K, Rangel-Moreno J, Nogales A, Martinez-Sobrido L. 2019. A Luciferase-fluorescent Reporter Influenza Virus for Live Imaging and Quantification of Viral Infection. J Vis Exp doi:10.3791/59890.

38. Nogales A, Ávila-Pérez G, Rangel-Moreno J, Chiem K, DeDiego ML, Martínez-Sobrido L. 2019. A novel fluorescent and bioluminescent Bi-Reporter influenza A virus (BIRFLU) to evaluate viral infections. J Virol doi:10.1128/JVI.00032-19.

39. Dietert K, Gutbier B, Wienhold SM, Reppe K, Jiang X, Yao L, Chaput C, Naujoks J, Brack M, Kupke A, Peteranderl C, Becker S, von Lachner C, Baal N, Slevogt H, Hocke AC, Witzenrath M, Opitz B, Herold S, Hackstein H, Sander LE, Suttorp N, Gruber AD. 2017. Spectrum of pathogen- and model-specific histopathologies in mouse models of acute pneumonia. PLoS One 12:e0188251.

40. Klopfleisch R. 2013. Multiparametric and semiquantitative scoring systems for the evaluation of mouse model histopathology--a systematic review. BMC Vet Res 9:123.

41. Zhao H, Doyle TC, Coquoz O, Kalish F, Rice BW, Contag CH. 2005. Emission spectra of bioluminescent reporters and interaction with mammalian tissue determine the sensitivity of detection in vivo. Journal of Biomedical Optics 10:41210.

42. Rodriguez JF, Rodriguez D, Rodriguez JR, McGowan EB, Esteban M. 1988. Expression of the firefly luciferase gene in vaccinia virus: a highly sensitive gene marker to follow virus dissemination in tissues of infected animals. Proc Natl Acad Sci U S A 85:1667–71.

43. Sánchez-Puig JM, Sánchez L, Roy G, Blasco R. 2004. Susceptibility of different leukocyte cell types to Vaccinia virus infection. Virol J 1:10.

44. Paley MA, Prescher JA. 2014. Bioluminescence: a versatile technique for imaging cellular and molecular features. Medchemcomm 5:255–267.

45. Tran V, Moser LA, Poole DS, Mehle A. 2013. Highly sensitive real-time in vivo imaging of an influenza reporter virus reveals dynamics of replication and spread. J Virol 87:13321–9.

46. De Baets S, Verhelst J, Van den Hoecke S, Smet A, Schotsaert M, Job ER, Roose K, Schepens B, Fiers W, Saelens X. 2015. A GFP expressing influenza A virus to report in vivo tropism and protection by a matrix protein 2 ectodomain-specific monoclonal antibody. PLoS One 10:e0121491.

47. Cheng BY, Ortiz-Riano E, de la Torre JC, Martinez-Sobrido L. 2015. Arenavirus Genome Rearrangement for the Development of Live Attenuated Vaccines. J Virol 89:7373–84.

48. Park JG, Ávila-Pérez G, Nogales A, Blanco-Lobo P, de la Torre JC, Martínez-Sobrido L. 2020. Identification and characterization of novel compounds with broad spectrum antiviral activity against influenza A and B viruses. J Virol doi:10.1128/JVI.02149-19.

